# Autophagy Cargo Profiles in Skeletal Muscle during Starvation and Exercise

**DOI:** 10.1101/2024.09.29.615610

**Authors:** Mohd Farhan, Shangze Lyu, Trezze P Nguyen, Dakai Zhang, Hong Liang, Yong Zhou, Yu A An, Hongyuan Yang, Guangwei Du, Yang Liu

## Abstract

Autophagy is a cellular process to clear unwanted and dysfunctional cellular cargoes, which are sequestered in autophagosomes before their delivery to lysosomes for degradation. Autophagy cargo selection, mediated by cargo receptors, varies across cell types and conditions. Understanding the cargo features is essential for elucidating autophagy’s function in specific physiological or pathological contexts. Here we present a simple and rapid method for isolating LC3B-positive autophagosomes from the tissues of GFP-LC3 transgenic mice, a widely used autophagy reporter model, without relying on the complex ultracentrifugation steps required by traditional methods. When combined with quantitative proteomics, this approach enables efficient in vivo characterization of autophagy cargoes. We applied this method to establish autophagy cargo profiles in skeletal muscle during starvation and exercise, two physiological conditions that activate autophagy, and identified distinct cargo selection patterns, with significantly higher levels of ER-phagy and ribophagy observed during starvation. We further revealed the ER-phagy receptors TEX264 and RETREG1/FAM134B as potential mediators of the elevated ER-phagy under starvation. In summary, we report an efficient workflow for in vivo autophagy cargo characterization and provide detailed analysis and comparison of cargo profiles under starvation and exercise conditions.

## Introduction

Macroautophagy/autophagy is an evolutionary conserved process to degrade unwanted and dysfunctional cytoplasmic components, known as autophagy cargoes. Autophagy plays a vital role in maintaining cellular homeostasis by removing harmful intracellular materials and facilitating the reuse of internal source of nutrients under stress conditions [1]. Autophagy cargoes are first engulfed by double-membrane vesicles called autophagosomes, which then deliver them to lysosomes for degradation. To date, numerous autophagy cargoes have been reported, including cytosolic proteins, protein aggregates, intracellular pathogens and nearly all the organelles [2]. As the crucial “cleaning” and “recycling” process in the cell, autophagy plays an essential role in maintaining normal cellular functions. And dysregulation of autophagy has been implicated in the pathogenesis of various diseases [3].

The initial step of autophagy involves the formation of an isolation membrane, called a phagophore, near the cytoplasmic cargo. The phagophore expands around the cargo and is ultimately sealed to form an autophagosome, with the cargo sequestered inside. The autophagosomes then fuse with lysosomes to form autolysosomes, where the cargo is degraded [4]. The initiation of autophagy requires the activation of ULK1/2 kinase, Class III phosphoinositide 3-kinase (PI3K), and serial downstream events mediated by many autophagy-related proteins [4]. A critical event during phagophore elongation is the conjugation of cytosolic ATG8 proteins to phosphatidylethanolamine (PE) on both sides of the phagophore membrane [4]. The lipidated ATG8 proteins play essential roles in phagophore elongation and closure as well as autophagy cargo capture, and remain on both outer and inner membranes of the autophagosomes to facilitate fusion with lysosomes [5]. After the formation of autolysosomes, ATG8 proteins on the cytosolic side of the membrane are cleaved off by ATG4 proteins [6].

Seven ATG8 proteins have been identified in human, categorized into two subfamilies: the LC3 family (Microtubule-associated proteins 1A/1B light chain 3; comprising LC3A, LC3B, LC3B2, and LC3C) and the GABARAP family (Gamma-aminobutyric acid receptor-associated protein; including GABARAP, GABARAPL1, and GABARAPL2) [7]. Although all ATG8 proteins are presumed to fulfill similar functions due to structural similarities, recent studies suggest that GABARAP family proteins may play more essential roles in autophagosome formation and fusion with lysosomes, while LC3 family proteins may be more important for cargo recognition [8]. However, the specific roles of different ATG8 proteins in distinct cell types and their potential functional redundancy require further clarification. Among the ATG8 protein family, LC3B is the most extensively studied for its roles in autophagosome biogenesis and turnover as well as cargo recognition [9]. Monitoring the lipidation of LC3B (hereafter referred to as LC3) is the most commonly employed method for assessing autophagy levels. For instance, a GFP-LC3B transgenic mouse strain (hereafter referred to as the GFP-LC3 mouse) was developed by N. Mizushima and colleagues to facilitate in vivo monitoring of autophagy [10], and has since been widely adopted by various research groups to investigate autophagy under diverse physiological and pathological conditions. In the cells of these transgenic mice, autophagosomes are decorated with lipidated GFP-LC3, and therefore can be visualized and quantified to reveal autophagy levels [10].

The autophagy process can be non-selective and/or selective. Non-selective autophagy involves the random engulfment of cytoplasmic material into autophagosomes for degradation, while selective autophagy targets specific cargoes [1]. The capture of these cargoes during selective autophagy is mediated by cargo receptors that bind to or reside on the cargoes. These receptors utilize a LC3-interacting region (LIR) to interact with lipidated ATG8 proteins on phagophores, facilitating the capture of the cargoes by phagophores and autophagosomes [11]. For instance, the cargo receptor SQSTM1/p62 binds to the ubiquitinated substrates and ATG8 proteins, directing these substrates to autophagosomes [12]. In well-studied forms of selective autophagy, such as mitophagy (the degradation of mitochondria) and ER-phagy (the degradation of the endoplasmic reticulum [ER]), multiple receptors have been identified that may function under distinct conditions and in different cell types [13,14].

As a defense mechanism against nutrient and energy stresses, autophagy is activated during both fasting (starvation) and endurance exercise—two conditions known to confer beneficial effects on health in humans and model organisms, such as slowing down the aging process and combating age-related diseases [15]. Notably, autophagy activation has been shown to contribute, at least partially, to these positive outcomes [16–18]. Despite the important role of autophagy during fasting and exercise, the nature of the cargoes in these two conditions remains unknown.

One primary function of autophagy is to degrade the cargoes, therefore, a thorough understanding of autophagy cargo properties under specific physiological or pathological conditions would provide mechanistic insights into how autophagy fulfills its roles. The rapid development of high-throughput omics technologies has enabled detailed characterization of organelles [19–21], yet our understanding of the features of autophagosomes in vivo, including their cargoes, remains incomplete due to the lack of effective isolation methods. Conventional techniques, such as density gradient ultracentrifugation, are complex and time-consuming, requiring large amounts of starting material, which limits their feasibility [22]. Here we introduce a simple, immunoaffinity-based method for isolating LC3B-positive autophagosomes from the tissues of GFP-LC3 mice using relatively small amounts of starting material. When combined with quantitative proteomics, this method allows the characterization of autophagy cargo in vivo. We apply this approach to establish autophagy cargo profiles in skeletal muscle during starvation and exercise, and reveal distinct cargo selection patterns under these two conditions. We anticipate that this method, along with the very first analysis and comparison of autophagy cargo features during starvation and exercise, will serve as a valuable resource for the field of autophagy research.

## Results

### Isolation of autophagosomes from the tissues of GFP-LC3 mice for downstream characterization

As a widely used autophagy reporter, the GFP-LC3 mouse strain allows for in vivo monitoring of autophagy levels [10]. GFP-LC3 is expressed at adequate levels across a variety of tissues in this mouse model [10]. The presence of lipidated GFP-LC3 on autophagosome membrane enables immunoaffinity-purification of autophagosomes using anti-GFP antibodies. Although several studies have employed this strategy [23,24], apparent caveats exist in these published methods, including the inability to assess non-specific binding of cytoplasmic components to the GFP antibody-coupled beads and visible contamination from unlipidated cytosolic GFP-LC3 in the preparations. To address these issues, we took the following approaches. First, when processing tissues from GFP-LC3 mice for autophagosome isolation, we included control tissues from wild-type (WT) mice without GFP-LC3 expression alongside. This allows us to determine the non-specific binding signals and accurately reflect the true signals coming from autophagosomes. Second, we performed size exclusion chromatography (SEC) on the tissue lysates to completely remove cytosolic GFP-LC3 (**Fig. 1A and B**). Due to their larger size, autophagosomes elute in the earlier fractions during SEC, as reflected by the presence of lipidated GFP-LC3 and LC3 (marked as II), together with other large cellular components. In contrast, the cytosolic GFP-LC3 and LC3 (marked as I) are only found in the later fractions (**Fig. 1B**). Notably, lipidated GFP-LC3 and LC3 are still present in these later fractions, suggesting the existence of smaller structures containing lipidated LC3, potentially phagophores or fragmented autophagosomes. Furthermore, we observed that ATG5-ATG12 conjugate and phosphorylated ATG16L1 (Ser278), markers for phagophores [25,26], are absent from the earlier fractions and present in the later ones, providing additional evidence suggesting that the earlier fractions contain mature autophagosomes while the later fractions contain phagophores (**Fig. 1B**). Thus, we collected the earlier fractions for autophagosome isolation using anti-GFP nanobody tandem-conjugated magnetic beads (see Materials and Methods for details). This workflow enabled effective purification of autophagosomes from the skeletal muscles of starved GFP-LC3 mice (both quadriceps muscles from one mouse used per isolation), as indicated by the higher total protein abundance in the isolation preparation from GFP-LC3 mice compared to that from WT control mice (**Fig. 1C**) as well as the enrichment of lipidated GFP-LC3, LC3 and SQSTM1/p62 in the preparation (**Fig. 1D**). The isolation efficiency was approximately 49.9± 2.6%, calculated by comparing the depletion of GFP-LC3 II after isolation to the total amount present before isolation (**Fig. 1D**). Moreover, we also observed the presence of the marker proteins from other cellular components including cytosol (GAPDH), mitochondria (COXIV), ER (SEC61A1), ribosome (40S ribosome protein S6), Golgi (GM130), plasma membrane (ATP1A2), early endosome (Rab5A), late endosome (Rab7) and lysosome (Cathepsin D), all of which have higher abundance in the GFP-LC3 samples than in the WT control samples. It is somewhat expected since many of these cellular parts (such as cytosol, mitochondria, ER and ribosome) are common autophagy cargos, while the presence of others proteins such as Rab5A, Rab7 and Cathepsin D may suggest the presence of newly-formed autolysosomes and/or CASM (Conjugation of ATG8 to Single Membrane) vesicles in the preparation since these structures also have GFP-LC3 on the membrane therefore can be pulled-down as well (See Discussion for the limitation of the method). Unlike GFP-LC3 or LC3 whose levels were significantly depleted from the input after the isolation procedure, the levels of the majority of these marker proteins, except GAPDH and Cathepsin D, did not significantly change, suggesting a specific enrichment of autophagosomes during isolation. Because most free cytosolic proteins, including GAPDH, were removed during the SEC process, the remaining GAPDH observed in the input was likely enclosed within vesicular structures—presumably autophagosomes. Consequently, a depletion of GAPDH was also observed following the isolation procedure (**Fig. 1D**). The enrichment of Cathepsin D is likely due to the presence of newly-formed autolysosomes, which can still retain LC3 on their membranes prior to its cleavage, allowing them to be co-purified during the process. We did not quantify p62 depletion after the isolation procedure due to the presence of high concentrations of BSA in the input, which was added to reduce non-specific binding and has a similar molecular weight to p62, making it difficult to detect p62 on the western blots. Furthermore, we performed a trypsin digestion assay and found that cargo protein p62, ER protein Calnexin (antibody recognizing the cytoplasmic, C-terminus of Calnexin) and S6 were protected from trypsin digestion (**Fig. 1E**). These results not only indicate that the isolated autophagic vesicles were sealed, but also confirm that the detection of cargo proteins such as Calnexin and S6 was not due to non-specific binding to the beads. Instead, their resistance to trypsin digestion demonstrates that these proteins were enclosed within the autophagosomes. Further analysis with cryo-electron microscope (cryo-EM) reveals double-membrane vesicles that resemble autophagosomes in the preparation (**Fig. 1F**). Notably, despite the concentration of vesicles on the cryo-EM grid was low because of the insufficient elution of vesicles from the beads, presumably due to the ultra-high affinity between GFP nanobody and GFP-LC3, out of the total 22 vesicles with good morphology, 16 of them (∼73%) have double-membraned structures. The ones with single membrane may be autolysosomes and/or CASM vesicles (see Discussion). Furthermore, to achieve more effective pull-down of autophagosomes within a short timeframe, we leveraged a newly developed anti-GFP nanobody tandem [27] expressed and purified from *E.coli* (**Fig. S1A**), which exhibits ultra-high affinity for GFP. Compared to single GFP nanobody, the use of the nanobody tandem resulted in a slightly more effective autophagosome isolation, as evidenced by higher levels of lipidated GFP-LC3 and LC3 as well as SQSTM1/p62 in the preparation (**Fig. S1B**).

**Figure 1.**
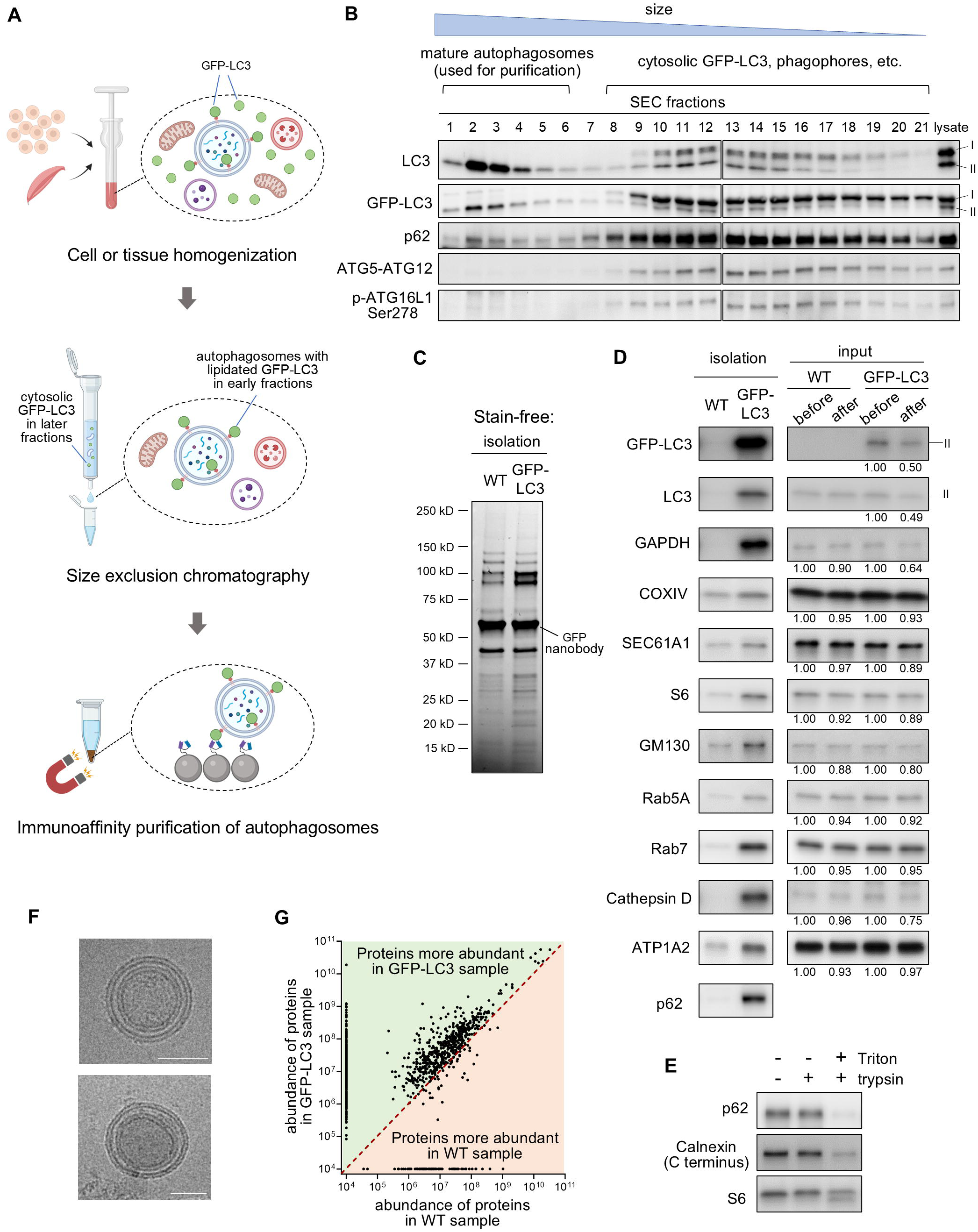
A novel method for isolating LC3B-positive autophagosomes from the tissues of GFP-LC3 mice. (**A**) Schematic illustration of the isolation process. Briefly, the tissues or cells are homogenized to release the cellular contents including cytosolic unlipidated GFP-LC3 (I) and autophagosomes containing lipidated GFP-LC3 (II). The lysates are then subjected to Size Exclusion Chromatography (SEC), during which process the large-sized autophagosomes are eluted earlier than the small-sized soluble GFP-LC3 I. Afterwards, the SEC fractions that contain autophagosomes are used for immunoaffinity-pull-down of autophagosomes using GFP nanobody-conjugated magnetic beads. (**B**) Western blots analysis shows the distribution of GFP-LC3 I and II, LC3 I and II, p62, ATG5-ATG12 conjugate and p-ATG16L1 Ser 278 in different SEC fractions of skeletal muscle lysate obtained from a starved GFP-LC3 mouse. The fraction numbers, ranging from 1 to 21, indicate the elution order from earliest to latest, corresponding to the largest to smallest sizes. Fractions 1 to 6 which contain mature autophagosomes are used for isolation. Similar results were observed in at least three independent experiments. (**C**) Stain-free method showing the amounts of total proteins in the isolation samples**. (D)** Left, western blots analysis shows the enrichment of autophagosome markers, GFP-LC3 II, LC3 II and p62 as well as markers of other cellular components in the autophagosome preparation from the skeletal muscles of a starved GFP-LC3 mouse compared to that from a WT control. GAPDH as a cytosol marker; COXIV as a mitochondria marker; SEC61A1 as an ER marker; S6 as a ribosome marker; GM130 as a Golgi marker; Rab5A as an early endosome marker; Rab7 as a late endosome marker; Cathepsin D as a lysosome marker; ATP1A2 as a plasma membrane marker. Similar results were observed in at least three independent experiments. On the right shows the levels of these proteins in the input (SEC eluates of WT and GFP-LC3 muscle lysates) before and after autophagosome isolation. The number underneath each lane indicates the abundance of the protein calculated from three experiments (the value before isolation was considered as 1). The isolation efficiency was determined by calculating the depletion of GFP-LC3 II after isolation relative to the total amount of GFP-LC3 II present before isolation based on the results from three independent experiments. The blots on the left were loaded with 1/12 of the isolated autophagosome samples, while the blots on the right used 1/500 of the total input. Both sets of western blots were processed in parallel under identical conditions. (**E**) Trypsin digestion assay on the autophagosome preparation from the muscles of a starved GFP-LC3 mouse in the presence or absence of 0.2% Triton X-100. (**F**) Cyro-EM images show the presence of autophagosomes with double membranes in the autophagosome preparations. Scale bar, 50 nm. (**G**) An example for using label-free quantitative proteomic to compare the abundance of proteins (expressed as spectrum intensity) identified in the autophagosome preparations from starved GFP-LC3 mice and WT mice. Individual dot represents a protein. Red dotted line indicates equal abundance in the two samples. Proteins locate next to Y and X axis are proteins only found in GFP-LC3 sample and WT sample, respectively.

To test the suitability of the purified autophagosomes for cargo characterization, we conducted label-free quantitative mass spectrometry analysis to compare protein abundance in the autophagosome preparations from the skeletal muscle of a starved GFP-LC3 mouse and a WT control mouse in a proof-of-concept experiment (**Fig. 1G**). We identified approximately 1,160 proteins with higher abundance in the GFP-LC3 sample, including 638 proteins unique to this sample. In comparison, we identified about 670 proteins in the WT sample, highlighting the importance of including such controls to account for non-specific binding signals (**Fig. 1G)**. If we define “proteins from autophagosomes” as those whose abundance in the GFP-LC3 sample is at least 1.5 times more than that in the WT sample—a threshold established in previous studies using similar immunoaffinity-based strategies for lysosome purification [28]—approximately 1,100 proteins meet this criterion, including dozens of known autophagosome proteins and autophagy receptors, namely LC3B, FYCO1 [29], SQSTM1/p62, NBR1 [30], RETREG1/FAM134B (Reticulophagy regulator 1) [31], TEX264 [32,33], BCL2L13 [34], PHB2 [35], *etc* (**Supplementary Dataset 1**). Thus, the autophagosome isolation method proposed here represents a powerful tool for the rapid purification (in ∼2.5 hours) of LC3B-positive autophagosomes from GFP-LC3 mouse tissues using relatively small amounts of starting material (e.g., the quadriceps muscles from one mouse under starvation condition), compared to previous studies that required livers from multiple mice for isolation [22], and without relying on the complex ultracentrifugation steps required by traditional methods.

### Autophagy cargo profiles during starvation and exercise in skeletal muscle

Fasting (starvation) and exercise are two major physiological conditions that activate autophagy, which helps recycle intracellular nutrients in response to the nutrient/energy stresses under these two conditions. Both starvation and exercise induce autophagy in various tissues of mice, including skeletal muscle, cardiac muscle, liver, pancreas, *etc* [10,17]. While autophagy activation induced by nutrient/energy stresses was initially thought to be non-selective, growing evidence suggests that during bulk autophagy, some cargoes (such as cytosolic proteins) are engulfed into autophagosomes in a non-selective way, while others are selectively targeted to autophagosomes through cargo receptors [11]. For example, during starvation in cell lines, ER and ribosomes can be specifically chosen for autophagic degradation through the actions of ER-phagy receptors and ribophagy receptor, respectively [28,32,33]. In addition, it has been shown that mitophagy is activated by exercise in skeletal muscle, although the responsible receptor(s) remain unclear [36]. Despite these understandings, a comprehensive analysis of autophagy cargoes in mouse tissues under starvation and exercise conditions is lacking. Therefore, we applied the workflow described above to address this knowledge gap.

We first assessed the levels of lipidated GFP-LC3 II in the autophagosome-containing SEC fractions of skeletal muscle lysates from mice subjected to starvation, exercise, or left unmanipulated (basal condition), which reflect the abundance of autophagosomes. As expected, both starvation and exercise significantly increased GFP-LC3 II levels compared to the basal condition, with approximately 1.9-fold and 1.8-fold increases, respectively (**Fig. S2A**). Based on these results, in order to isolate equivalent amounts of LC3 II positive autophagosomes, we used the quadriceps muscles from one mouse for the starvation or exercise condition, while combining the quadriceps muscles from two mice for the isolation in basal condition. Compared to those from WT control mice, the autophagosomes preparations from GFP-LC3 mice in all three conditions showed enrichment of GFP-LC3 II, LC3 II and SQSTM1/p62 (**Fig. S2B**).

We then performed label-free quantitative proteomics to analyze the amounts of proteins identified in the autophagosome preparations from the three conditions. We conducted four biological replicates for each condition, and we defined proteins as “from autophagosomes” in each replicate based on a criterion that the protein’s abundance in the GFP-LC3 sample is at least 1.5-fold greater than in the corresponding WT control sample (**Supplementary Dataset 1**). We then sought the common proteins identified in all four replicates of one condition that are significantly more abundant in the GFP-LC3 samples compared to WT control, and found 272 proteins in basal condition, 515 proteins in starvation condition, and 365 proteins in exercise condition (**Fig. 2A; Supplementary Dataset 1**). To establish the autophagy cargo profiles in each condition, we next analyzed the cellular components to which these proteins belong using subcellular localization information from the Uniprot database **(Supplementary Dataset 1**). Next, we employed two analytical approaches to reflect autophagy cargo selection patterns under different conditions. In the first analysis, we calculated the total number of proteins from different cytoplasmic components-including mitochondria, cytosol, plasma, membrane, nucleus, ER (or SR [Sarcoplasmic Reticulum] in muscle), Golgi, ribosomes, peroxisomes and lipid droplets-and determined the percentage of proteins from each cytoplasmic component relative to the total number of proteins in each condition (**Fig. 2B; Supplementary Dataset 1;** proteins that are localized to multiple components are counted for each of their respective localizations). We excluded endosome, transport vesicle and lysosome from this analysis, as it is challenging to tell whether the proteins localized to these compartments are autophagy cargos or merely present on the membranes of autolysosomes and other GFP-LC3 II-positive vesicles, which may be co-isolated in the autophagosome preparation (see Discussion). We found that there are both larger number and higher percentage of proteins from mitochondria or peroxisome in the autophagosomes from exercise condition compared to the control or starvation condition, and larger number and higher percentage of proteins from plasma membrane, ER and ribosome in the autophagosomes from starvation condition compared to the control or exercise condition (**Fig. 2B)**. In the second analysis, we calculated the total abundance of proteins (normalized to the abundance of LC3B in each sample) from different cellular components (**Fig. 2C; Supplementary Dataset 1;** proteins that are localized to multiple components are counted for each of their respective localizations). Assuming that the same number and set of proteins from a specific cellular part was identified in the autophagosomes from two different conditions, if the abundance of these proteins is higher in one condition than the other, this still suggests higher amounts of the specific cellular part are targeted for autophagic degradation in this condition. Thus, we believe that this analysis method, which calculates not only the number but the abundance of proteins from different cellular components in the autophagosome preparations, could accurately reflect on the autophagy cargo selection patterns in different conditions. We found that in both basal and exercise conditions, proteins from mitochondria are most abundant in autophagosomes, followed by proteins from cytosol and plasma membrane (**Fig. 2C**). In the exercise condition, there was a significant increase in the abundance of proteins from the plasma membrane and ER compared to the basal condition. There was also a trend indicating higher abundance of peroxisomal and nuclear proteins in the exercise condition compared to the basal condition (*p*=0.051 and *p*=0.071, respectively) (**Fig. 2C**). In addition, while there were larger number of mitochondrial proteins identified in the exercise condition than in the basal condition (**Fig. 2B**), no significant increase in the abundance of these proteins was observed, despite a trend toward higher (*p=*0.15; **Fig. 2C**). A possible explanation is that mitophagy is most active during the recovery period post-endurance exercise, not immediately after exercise [37]. In starvation condition, ER (SR) proteins are the most abundant in autophagosomes, with levels drastically higher than those in both basal and exercise conditions (**Fig. 2C**). Among the SR proteins identified in the autophagosomes from starvation condition, several Ca^2+^ handling molecules, namely SERCAs (Sarcoplasmic/endoplasmic reticulum calcium ATPase 1 to 3) and Calsequestrin-1 were highly abundant (together constituting approximately 30% of the total protein abundance in the autophagosome preparations), and are absent in the autophagosomes from basal and exercise conditions (**Fig. S3; Supplementary Dataset 1**). This suggests a potential preference for targeting these molecules or ER subdomains containing them for autophagic degradation during starvation in skeletal muscle. Given the extreme high abundance of these SR proteins, potential false-positive identification of them in starvation condition or false-negative identification in basal or exercise conditions would greatly impact the analysis results. To mitigate the risk, we also analyzed the abundance of other ER proteins, excluding SERCAs and Calsequestrin-1, in the autophagosomes from starvation condition, and found that their levels remained significantly higher than those in autophagosomes from basal or exercise condition (**Fig. S3**). Based on these results, we concluded that autophagosomes in starvation condition contained a greater abundance of ER proteins compared to those in basal or exercise condition. Additionally, we observed a significantly higher abundance of ribosomal proteins in the autophagosomes from starvation conditions compared to both basal and exercise conditions in which no ribosomal proteins were identified (**Fig. 2B and C**). Autophagosomes from starvation condition also contained significantly higher abundance of proteins from cytosol, plasma membrane and nucleus than those from basal condition, while showing a significantly lower abundance of peroxisomal proteins compared to those from exercise condition (**Fig. 2C**). Notably, there was also a trend suggesting that the abundance of mitochondrial proteins was lower in autophagosomes from starvation compared to those from exercise condition (*p*=0.09; **Fig. 2C**). Based on our findings from the two different analyses comparing starvation and exercise conditions, we conclude that autophagosomes from the starvation condition contain dramatically higher amounts of ER and ribosomes compared to those from the exercise condition. In contrast, the exercise condition showed significantly more proteins from peroxisomes and a trend toward more mitochondrial proteins in the autophagosomes compared to the starvation condition, although these differences were not as pronounced as those seen for ER and ribosomal proteins.

**Figure 2.**
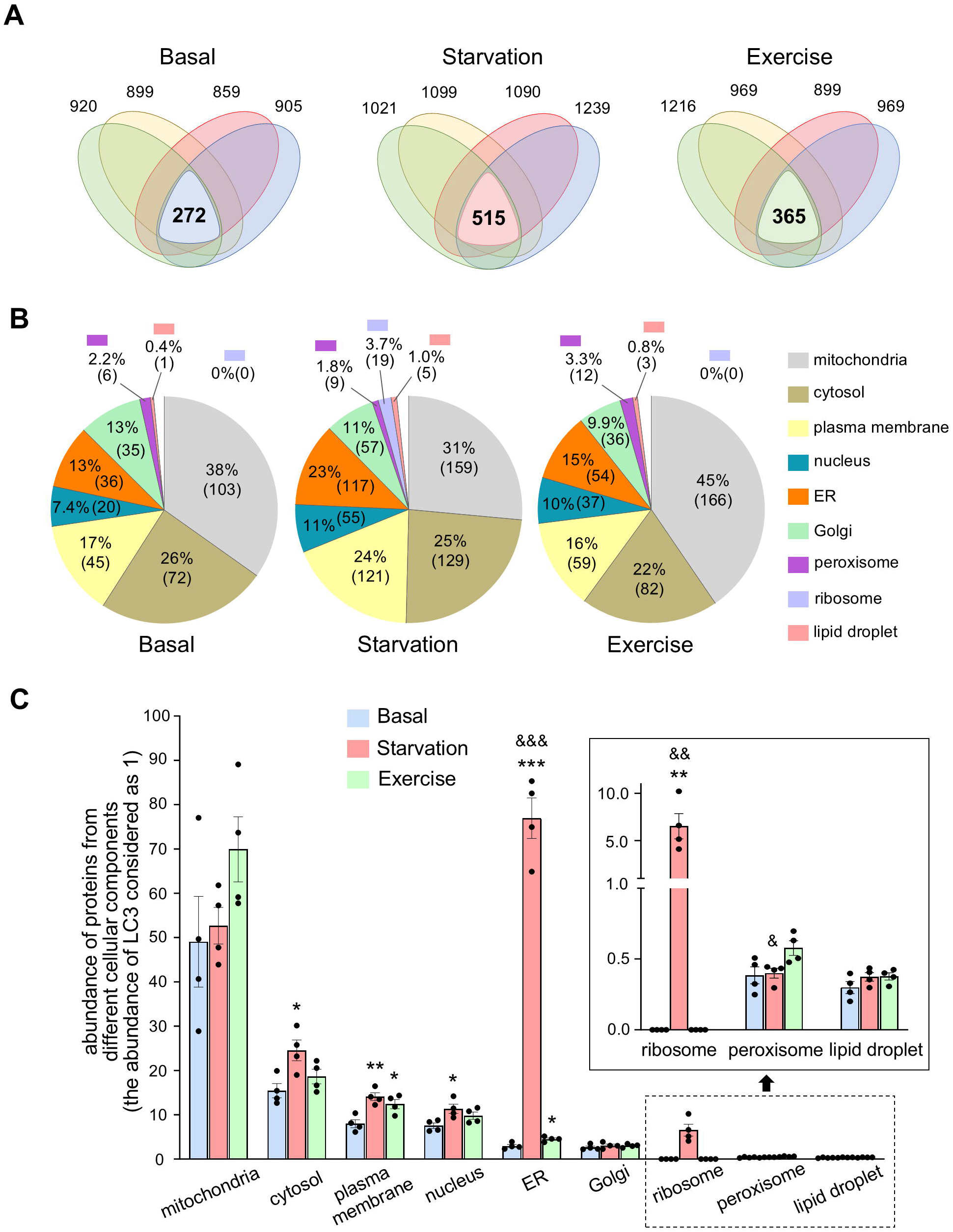
Autophagy cargo profiles in skeletal muscle during starvation and exercise. (**A**) The identification of proteins in the autophagosome preparations from skeletal muscle under basal, starvation and exercise conditions. In each biological replicate for each condition, proteins that are at least 1.5 times more abundant in the autophagosome preparations from the GFP-LC3 muscles than the corresponding WT control muscles are considered as “from autophagosomes. Each oval stands for one replicate, and the number near it indicates how many proteins are identified as “from autophagosomes” in this replicate. The number in the overlapping region indicates the common proteins identified in all four replicates that are significantly more abundant in GFP-LC3 samples compared to WT samples (*p* ≤ 0.05). These proteins were subjected to further analysis in (B) and (C). (**B**) The modified pie charts illustrate the percentage of proteins from a specific cytoplasmic component relative to the total number of proteins identified in the isolated autophagosomes under basal, starvation and exercise conditions. The number in the parentheses indicates how many proteins are from the specific cytoplasmic component. Proteins that are localized to multiple components were counted for each of their respective localizations. Thus, the sum of the percentages of all cytoplasmic components is more than 100%, and the pie charts are only approximate to reflect the portions. Refer to the values on the chart for accurate assessment. (**C**) The total abundance of proteins from various cellular components identified in the isolated autophagosomes under basal, starvation and exercise conditions. The abundance of proteins is normalized to the abundance of LC3B in each replicate. Proteins that are localized to multiple components are counted for each of their respective localizations. * *p* < 0.05, ** *p* <0.01, *** *p* < 0.001 when compared to basal condition. $ *p* < 0.05, $$ *p* < 0.01, $$$ *p* < 0.001 when compared to exercise condition. Two sides, unpaired Student’s t-test was used for statistical analysis. Data are mean ± s.e.m.

We next investigated whether the dramatically higher abundance of ER and ribosomal proteins in autophagosomes from starvation condition was simply a result of more ER and ribosomes available for autophagic degradation in the muscle cells during starvation. To address this, we performed label-free quantitative proteomics to analyze the abundance of proteins in the total tissue lysates from all three conditions, using three replicates for each condition. We identified common proteins across all replicates and found 474 proteins in basal condition, 471 proteins in starvation condition, and 493 proteins in exercise condition (**Fig. S4A; Supplementary Dataset 2**). Next, using these common proteins, we calculated the abundance of proteins from different cytoplasmic parts (total protein abundance considered as 100 in each sample; **Fig. S4B; Supplementary Dataset 2**). No significant differences in the abundance of proteins from ER and ribosomes were observed across the three conditions (**Fig. S4B)**. These results suggested that the higher abundance of ER or ribosomal proteins in the autophagosomes from starvation condition is likely due to enhanced selective targeting of these components for autophagic degradation. In summary, we analyzed the autophagy cargo profiles in basal, starvation and exercise conditions in mouse skeletal muscle, and revealed distinct autophagy cargo selection patterns with dramatically higher levels of ER-phagy and ribophagy during starvation. To our knowledge, this study is the first to thoroughly examine and compare autophagy cargos in vivo during starvation and exercise, and the results suggested that even though both conditions induce general autophagy (bulk autophagy) in response to the nutrient/energy stress, there are certain differences in cargo selection patterns between the two.

### Assessment of mitophagy and ribophagy in basal, starvation and exercise conditions

To verify the autophagy cargo profiles obtained using the autophagosome isolation-proteomic workflow (**Fig.2**), we specifically examined mitophagy and ribophagy under basal, starvation and exercise conditions. Mitophagy was assessed in Mito-QC transgenic mice which express a functional inert, tandem mCherry-GFP protein fused to the mitochondrial targeting sequence of the outer mitochondrial membrane protein, FIS1 [38]. Under steady-state conditions, the mitochondria decorated with this overexpressed protein shows both red and green fluorescence. Upon mitophagy, mitochondria are delivered to lysosomes where mCherry fluorescence remains stable, but GFP fluorescence becomes quenched by the acidic microenvironment. This results in punctate mCherry-only foci that can be easily quantified as an indication of mitophagy levels (mitophagy flux). Consistent with previous findings [38], we observed mitochondrial reticulum within the muscle fiber, especially with the GFP channel. Though the signal from the mCherry channel on these reticulum structures was relatively faint, the mCherry-only puncta were obvious (**Fig. 3A**). In addition, although red-only puncta were observed in muscle from mice under all tested conditions, their number was significantly higher under the exercise condition compared to basal or starvation (**Fig. 3A** and **B**), suggesting increased mitophagy flux during exercise. This result is consistent with the cargo profiling data, which revealed a trend toward higher abundance of mitochondrial proteins in autophagosomes isolated under exercise condition (**Fig. 2C**). We also assessed ribophagy levels in skeletal muscle under basal, starvation, and exercise conditions by examining the colocalization of the ribosomal protein S6 with the lysosomal marker LAMP1, which reflects the delivery of ribosomes to lysosomes (ribophagy flux). Consistent with the cargo profiling data, we observed significantly more such colocalization under starvation condition compared to basal or exercise condition, suggesting higher ribophagy (**Fig. 3C, D**). These findings further demonstrate the utility of the autophagosome isolation–proteomics workflow as a robust method for profiling autophagy cargo in vivo.

**Figure 3.**
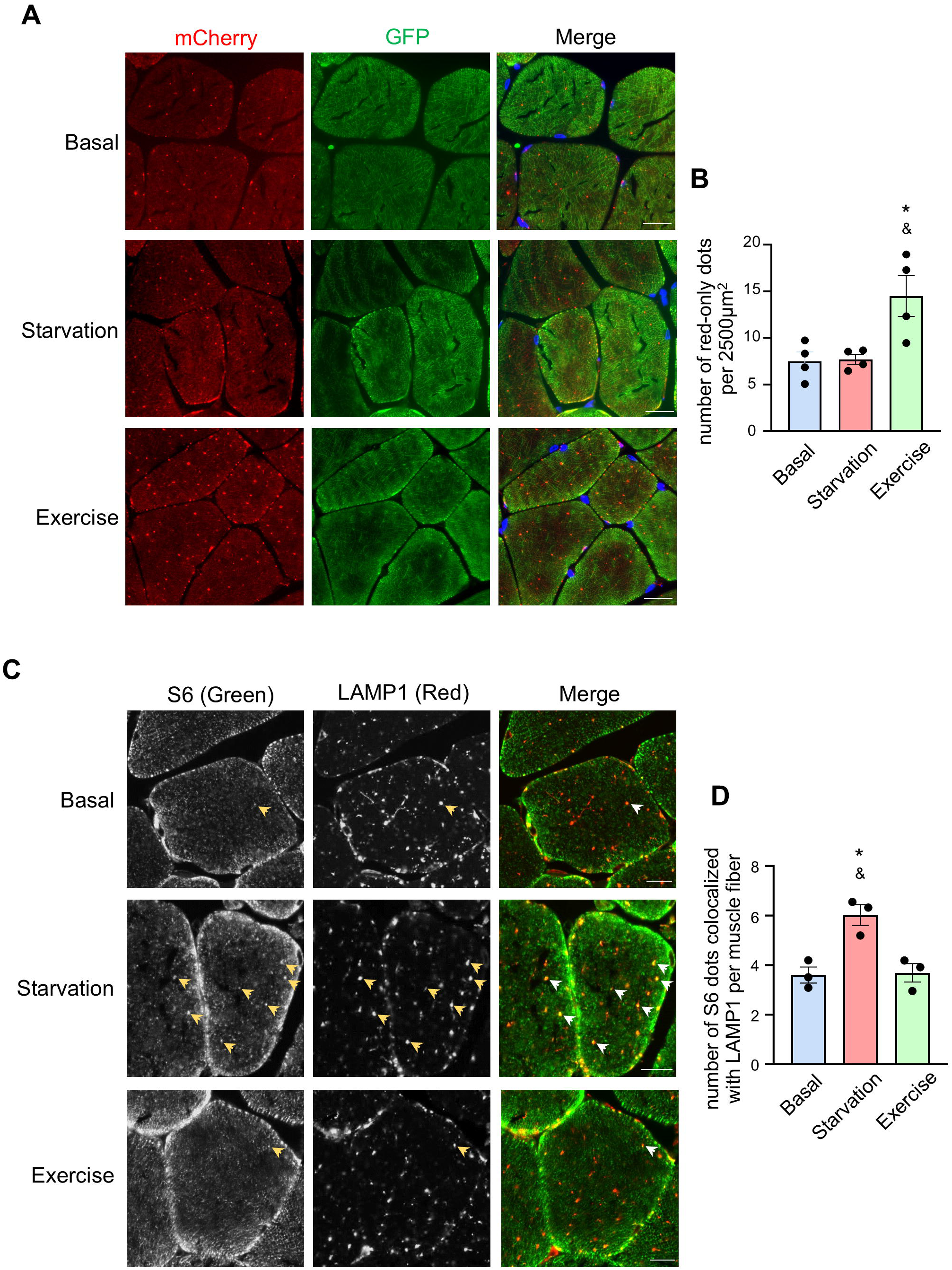
Mitophagy and ribophagy in skeletal muscles under basal, starvation and exercise conditions. **(A) (B)** Mitophagy levels in basal, starvation and exercise conditions assessed in the muscles of Mito-QC mice. The numbers of red-only dots reflect the mitophagy flux and were quantified in **(B).** Data points are individual mice (n = 4). * *p* <0.05 using two sides, unpaired Student’s t-test when comparing with basal condition. & *p* <0.05 using two sides, unpaired Student’s t-test when comparing with starvation condition. Data are mean ± s.e.m. Scale bar, 20 μm. **(C) (D)** Ribophagy levels in basal, starvation and exercise conditions assessed by examining the colocalization of ribosome protein S6 with lysosome protein LAMP1 in the muscles of wild-type mice. The numbers of colocalization dots per muscle fiber were quantified in **(D).** Data points are individual mice (n = 3), and 40 to 60 fibers were quantified for each mouse. * *p* <0.05 using two sides, unpaired Student’s t-test when comparing with basal condition. & *p* <0.05 using two sides, unpaired Student’s t-test when comparing with exercise condition. Data are mean ± s.e.m. Scale bar, 10 μm. Arrowheads denote the colocalization dots.

### Potential contribution of RETREG1/FAM134B and TEX264 to the elevated ER-phagy during starvation

Given the dramatic differences in ER-phagy observed between starvation and exercise conditions, we conducted additional experiments to investigate the underlying mechanisms and the potential cargo receptors involved. To date, several ER-phagy receptors have been discovered in mammalian cells, including RETREG1/FAM134B, TEX264, SEC62, RTN3L, CCPG1, *etc* [13]. To gain further insights into the potential mechanisms of the differential regulation of ER-phagy during starvation and exercise, we checked the presence of ER-phagy receptors in the autophagosome preparations from both conditions. We found that RETREG1/FAM134B, TEX264 and RTN3 are present in the autophagosomes from starvation conditions but not in those from exercise condition (**Supplementary Dataset 1**). Given this distinction, we chose to further explore the potential roles of RETREG1/FAM134B and TEX264 in the differential regulation of ER-phagy during starvation and exercise. We opted not to pursue RTN3 because only the full-length form of RTN3, RTN3L, contains the LIR domains and functions as an ER-phagy receptor [39], however, this isoform has very weak expression in skeletal muscle [40]. Indeed, none of the RTN3 peptides identified in our Mass spectrometry experiments are mapped to the specific region of RTN3L (**Table S1**). To confirm the functional roles of RETREG1/FAM134B and TEX264 as ER-phagy receptors contributing to enhanced ER-phagy during starvation, we then examined their interaction with lipidated GFP-LC3 II under starvation and exercise conditions. Using co-immunoprecipitation assays, we observed that RETREG1/FAM134B interacted with GFP-LC3 II and the binding was significantly more in starvation condition than in exercise condition (**Fig. 4A and B**). Notably, we consistently observed higher levels of RETREG1/FAM134B in the input from the starvation condition compared to exercise. This difference may reflect transcriptional or translational changes, or differential localization of RETREG1/FAM134B to ER subdomains that vary in solubility during lysate preparation. Therefore, we propose that total RETREG1/FAM134B levels may not accurately reflect ER-phagy activity. Instead, the increased association of RETREG1/FAM134B with LC3B under starvation provides stronger evidence for its role in regulating ER-phagy in this context. Similarly, we detected a markedly increased binding of GFP-LC3 II to TEX264 during starvation versus exercise (**Fig. 4C and D).** These results indicate that both RETREG1/FAM134B and TEX264 are likely to contribute to the elevated levels of ER-phagy observed in skeletal muscle during starvation.

**Figure 4.**
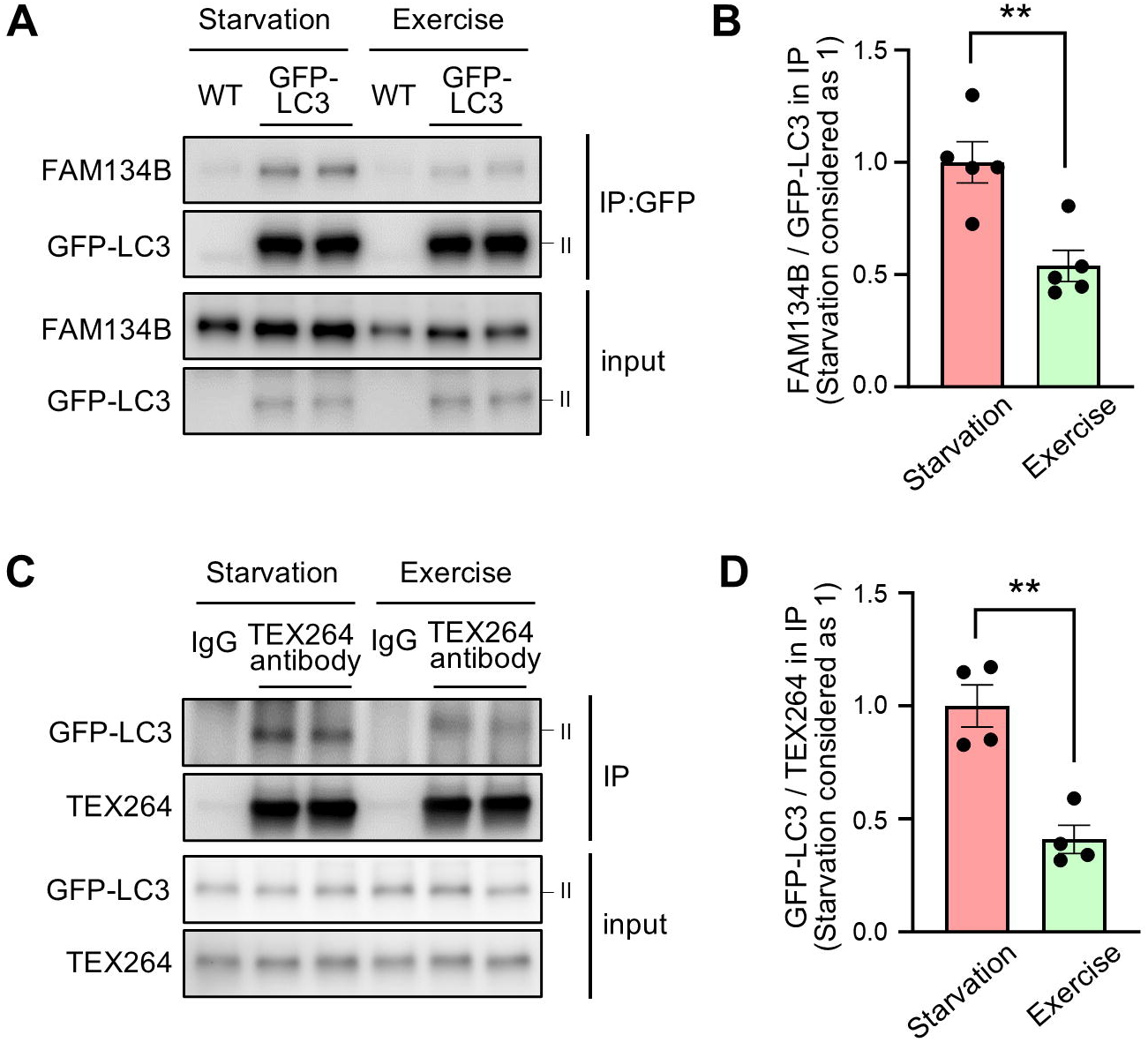
Potential contribution of RETREG1/FAM134B and TEX264 to the elevated ER-phagy during starvation. (**A**) (**B**) Co-immunoprecipitation of FAM134B with GFP-LC3 II in skeletal muscle lysates from WT and GFP-LC3 mice (post SEC to eliminate GFP-LC3 I) shows more interaction between these two molecules under starvation condition than exercise condition. (A) representative blots. (B) quantification of the results from two independent experiments. Data points are individual mice (n = 5). (**C**) (**D**) Co-immunoprecipitation of GFP-LC3 II with TEX264 in skeletal muscles lysates from GFP-LC3 mice (post SEC to eliminate GFP-LC3 I) shows more interaction between these two molecules under starvation condition than exercise condition. (C) representative blots. (D) quantification of the results from two independent experiments. Data points are individual mice (n = 4). ** *p* <0.01 using two sides, unpaired Student’s t-test. Data are mean ± s.e.m.

## Discussion

In this study, we characterized the properties of autophagy cargo in skeletal muscles during starvation and exercise, the two physiological conditions that activate autophagy, using a simple and rapid method for isolating LC3B-positive autophagosomes from the tissues of GFP-LC3 transgenic mice. We found that despite both conditions induce nutrient/energy stress, the cargo selection patterns under these two conditions are rather different, with significantly higher levels of ER-phagy and ribophagy during starvation compared to exercise.

One advantage of the autophagosome isolation method used in this study is that it does not require ultracentrifugation steps, which are used in most previously described methods [22,41]. The use of SEC enables the complete removal of cytosolic GFP-LC3 from tissue or cell lysates in less than 30 minutes (assuming the SEC columns are prepped when lysates are ready; see Materials and Methods). To ensure accuracy, a control sample from tissues or cells that do not express GFP-LC3 is processed in parallel to assess signals from proteins that might bind nonspecifically to the beads. Including this control is crucial, as we observed over 600 proteins binding nonspecifically in the control sample in a proof-of-concept experiment, with many detected exclusively in this sample (**Fig. 1E**). Comparing the protein abundance in the isolation sample to the control helps distinguish true autophagosome signals.

One limitation of the described autophagosome isolation method is that other lipidated GFP-LC3-positive vesicles, such as CASM (conjugation of ATG8 to single membranes)-related vesicles [42], may also be co-isolated with autophagosomes. Our observations that Rab5A and Rab7 were present in the isolation preparations and that single membrane vesicles were also observed in the preparation as revealed by cryo-EM, may reflect this limitation. A more rigorous control would involve performing the isolation in tissues with impaired autophagosome formation but intact LC3 conjugation to other membranes, such as FIP200-deficient tissues [43]. However, achieving these conditions in mice poses significant challenges. To reduce the influence of these additional vesicles on autophagy cargo characterization, we focused the cargo analysis on intracellular components such as various organelles while excluding endosomes and transport vesicles from our analysis. Using this approach, we anticipated minimizing the impacts of other LC3-positive vesicles on the cargo selection patterns. Another limitation of the isolation method is that it exclusively purifies GFP-LC3B-positive autophagosomes, potentially missing autophagosome populations that are negative for GFP-LC3B but positive for other mammalian ATG8 proteins. It remains unclear whether there are distinct subpopulations of autophagosomes that contain only specific ATG8 proteins while excluding others, or if autophagosomes generally incorporate all available ATG8 proteins. A recent study suggested that, at least in certain cell lines, more than 90% of autophagic vesicles contain both LC3B and GABARAP family proteins [41]. However, further investigation is needed to determine whether the entire autophagosome population in skeletal muscle is indeed LC3B-positive. This would help establish whether the autophagy cargo profiles obtained in this study provide a truly comprehensive view. Our approach used SEC to separate the cytosolic GFP-LC3 from the autophagosome-localized, lipidated GFP-LC3. However, although most of the lipidated GFP-LC3 or LC3 were present in the earlier fractions as expected, a small portion was detected in the later fractions. Since these later fractions contain ATG5–ATG12 and phosphorylated ATG16L1 (Ser278), we theorized that they may represent phagophores, which are smaller in size but still positive for lipidated GFP-LC3 or LC3. To this end, we cannot rule out the possibility that these later fractions also contain very small autophagosomes, and that our current approach may overlook the cargos within them. Further investigation is needed to explore this possibility.

In this study, we focused on skeletal muscle because a dramatic increase in autophagy levels following moderate-intensity aerobic exercise was observed in this tissue [17,44] and relatively large amount of material can be obtained from a single mouse. For researchers interested in applying the described workflow to isolate autophagosomes from tissues other than skeletal muscle, the required amount of starting material will need optimization. The quantity should be sufficient to yield enough autophagosomes for characterization without being excessive, as larger amounts may lead to increased non-specific binding, as evidenced in the WT control samples.

To obtain reliable autophagy cargo profiles under basal, starvation, and exercise conditions, we performed autophagosome isolation–proteomics on four biological replicates per condition. We noted a low number of identical proteins across replicates, which could reflect mouse-to-mouse variability or additional variation introduced during the experimental procedure. We observed significantly higher abundance of ER proteins and ribosome proteins in the autophagosomes from the starvation condition compared to those from the exercise condition. Since ER-bound ribosomes, which may account for up to 50% of total ribosomes [45,46], can be degraded along with ER via autophagy [32], and considering that the only known ribophagy receptor, NUFIP1 [28], was not identified in the autophagosome preparations from starvation condition, it is possible that the increased ribophagy observed during starvation is at least partially attributable to the enhanced ER-phagy. It is also possible that the increased ribophagy occurs independently of the enhanced ER-phagy, potentially mediated by an unidentified receptor. Further investigation is necessary to explore these possibilities. Our results suggested that two ER-phagy receptors, TEX264 and RETREG1/FAM134B, may contribute to the enhanced ER-phagy during starvation. This finding aligns with the results of previous studies using cell line models demonstrating that TEX264 is the major ER-phagy receptor which contributes to at least 50% of the total ER-phagy during starvation [32,33], and that RETREG1/FAM134B also contributes to starvation-induced ER-phagy [32]. We recognize that the involvement of different ER-phagy receptors during starvation may vary across different tissues, which requires further clarification in future studies.

The established autophagy cargo profiles during starvation and exercise in this study could shed light on the effects of intermittent fasting and endurance exercise, the two popular interventions for weight loss and fitness maintenance, on skeletal muscle physiology. For example, it has been shown that a high-fat diet induces ER stress in the skeletal muscle in mice, which is associated with and presumably contributes to insulin resistance [47,48]. Since ER-phagy can alleviate ER stress [49,50], the pronounced ER-phagy observed during starvation in muscle cells may contribute to improved insulin sensitivity in skeletal muscle, a well-documented benefit of intermittent fasting. Additionally, the increased mitophagy flux under exercise condition, consistent with previous reports, may help explain how exercise promotes the removal of damaged mitochondria, supports mitochondrial fitness and fuel efficiency, and ultimately enhances muscle performance [36,51].

In summary, in this study we present a novel and simple method for isolating LC3B-positive autophagosomes from the tissues of GFP-LC3 transgenic mice, enabling in vivo characterization of autophagosome properties, including cargo features. This method allowed us to uncover distinct autophagy cargo selection patterns in skeletal muscle during starvation and exercise, underscoring the complexity of autophagy regulation in response to various stressors.

## Materials and Methods

### Mouse strains

All mice were housed in the Center for Laboratory Animal Medicine and Care (CLAMC) at the University of Texas Health Science Center at Houston under 12-h light–dark cycles with ad libitum access to food and water. All the experimental procedures related to the mouse were approved by the institutional Animal Welfare Committee (AWC) and performed under institutional guidelines (protocol numbers: 21-0107 and 24-0105). Twelve- to sixteen-week-old male mice were used. GFP-LC3B transgenic mouse strain [10] was a kind gift from Dr. Noboru Mizushima and has been backcrossed with C57BL/6J mice for at least five generations. Wild-type (WT) and GFP-LC3B mice (heterozygous for GFP-LCB transgene) used in the experiments were littermates generated from the breeding between WT and heterozygous GFP-LC3B mice. Mito-QC transgenic mouse strain [38] was a kind gift from Dr. Ian Ganley, and heterozygous mice with Mito-QC transgene were used for experiments.

### Plasmids

Plasmid that expresses single GFP nanobody (GBP1/Enhancer) was a gift from Dr. Rebecca Voorhees (Addgene plasmid # 149336; http://n2t.net/addgene:149336; RRID: Addgene_149336). A tandem GFP nanobody (GBP1/Enhancer+ LaG16) was described in a previous study [27]. Plasmid that contains tandem GFP nanobody sequence was a gift from Dr. Motoyuki Hattori (Addgene plasmid # 140442; http://n2t.net/addgene:140442; RRID: Addgene_140442). The plasmid that expresses tandem GFP nanobody (GBP1/Enhancer+ LaG16) was generated by replacing the GBP1/Enhancer sequence in the single nanobody vector with the tandem nanobody sequence using Gibson assembly. Notably, sequences expressing the His14 tag, an Avi tag and a SUMO^EU1^ tag (can be cleaved by a SENP^EuB^ protease) are present in front of the nanobody sequence (discussed in the section “Preparation of GFP nanobody-conjugated magnetic beads” below).

### Starvation and exercise in mice

For starvation condition, mice were put into a new cage without food with water around 4:00 pm the day before experiment, and maintained in the cage for 18 hours (till 10:00 AM next day) before being euthanized for tissue collection. For basal condition, after the mice underwent an 18-hours starvation as described above, food was added back to the cage to refeed the mice for 4 to 5 hours before being euthanized for tissue collection.

For exercise condition, treadmill running in mice were performed as previously described [44] with minor modifications. In brief, mice were acclimated to the 10° uphill treadmill for two days. On day 1, mice ran for 5 min at a speed of 8 m/min followed by 2 min at a speed of 10 m/min. On day 2, mice ran for 5 min at a speed of 10 m/min followed by 2 min at a speed of 12 m/min. On day 3 and 4, mice were allowed to rest. Around 4 :00 pm on day 4, food was removed from the cages and the mice were starved for 18 hours. In the morning of day 5, food was put back into the cages to refeed the mice for 4 hours before the exercise experiment. Mice were subjected to a single bout of running starting at a speed of 10 m/min for 40 min. Afterwards, the speed was increased at a rate of 1 m/min every 10 min for a total of 30 min and then at a rate of 1 m/min every 5 min until finishing 18 m/min (total 95 min) when the mice were euthanized, and tissues were collected.

### GFP nanobody purification

The single or tandem GFP nanobody protein was expressed in NEBExpress^®^ *I^q^ E. coli* (New England Biolab, C3037I) as described previously [52]. Briefly, an 80 ml pre-culture of the *E. coli* containing the expression plasmid was grown in Super Broth (Research Products International, S33050) overnight at 28°C and diluted to 800 ml Super Broth the next morning. After dilution, the culture was incubated at 18°C until reaching OD600 value around 0.9 (normally in 4 to 5 hours). The expression of the protein was then induced by addition of 0.2 mM IPTG for 18 to 20 hours at 18°C. Cells were harvested by centrifugation and the pellet was resuspended in 70 ml lysis buffer (20 mM Hepes pH 7.5, 300 mM NaCl, 20 mM imidazole, 1 mM DTT, 1 mM PMSF) and lysed by a French press. The lysate was cleared by centrifugation for 45 min at ∼35,000 *g* at 4°C. The supernatant was then incubated with 1.5 ml bed volume Ni^2+^-NTA resin (Thermo Scientific, 88221), which has been equilibrated to lysis buffer, for 1 hour at 4°C before transferred to a gravity flow column. The resin was then sequentially washed with three column volume (CV) lysis buffer, 10 CV wash buffer A (20 mM Hepes pH 7.5, 300 mM NaCl, 50 mM imidazole, 1 mM DTT), 10 CV wash buffer B (20 mM Hepes pH 7.5, 300 mM NaCl, 100 mM imidazole, 1 mM DTT), and then eluted with elution buffer (20 mM Hepes pH 7.5, 300 mM NaCl, 500 mM imidazole, 1 mM DTT, 250 mM sucrose). Fourteen fractions of elution with 0.5 ml per fraction were collected and the presence of GFP nanobody in each fraction was determined by SDS-PAGE using the Mini-PROTEAN TGX stain-free precast gels (Bio-Rad, 4568096). The five fractions with the highest concentration of GFP nanobody were combined (final volume around 2.5 ml), and the buffer was then exchanged to storage buffer (20 mM Hepes pH 7.5, 200 mM NaCl, 1 mM DTT and 250 mM sucrose) using a PD-10 desalting column (GE Healthcare, 17085101) according to the user instructions. After the final protein concentration measurement, the purified nanobody was aliquoted, snap-frozen in liquid nitrogen, and stored at −80°C until further use.

### Preparation of GFP nanobody-conjugated magnetic beads

Before autophagosome isolation experiments were performed, GFP nanobody protein was conjugated to NHS-activated magnetic beads (Thermo Scientific, 88827). For each 100 µl beads, 120 µg tandem GFP nanobody (or 80 µg single nanobody) was used for conjugation. The conjugation steps were performed according to the user guide of the beads except that different coupling buffer (20 mM Hepes pH 7.5, 200 mM NaCl) and storage buffer (KPBS [potassium phosphate-buffered saline] with 2 µg/µl BSA) were used. The prepared beads can be stored at 4°C for up to a week before usage.

Notably, the GFP nanobody used in this study has three tags at the N terminus, a His14 tag for purification, an Avi tag that can be biotinylated by the biotin ligase BirA, and a SUMO^EU1^ tag that can be cleaved by a SENP^EuB^ proteinase. We tried several strategies for nanobody/beads coupling before we decided to use the current one. We first tried to biotinylate the Avi tag and couple the nanobody to streptavidin beads. However, although the coupling of nanobody to beads was very effective, we found high levels of proteins nonspecifically bound to the prepared beads during the autophagosome isolation step. We also tried using SENP^EuB^ proteinase (which cut SUMO^EU1^) to cleave off the nanobody together with the bound autophagosomes to reduce the non-specific binding, however, the efficiency of releasing autophagosomes from the beads was very low, possibly due to that each autophagosome has many GFP-LC3 molecules that can bind many GFP-nanobody molecules on the beads. We noticed that when GFP nanobody protein was coupled to NHS-activated beads as described above, the non-specific binding of proteins to the beads during autophagosome isolation step was much lower compared to the streptavidin beads. The conjugation of protein to the NHS beads relies on the formation of stable amide linkages between the primary amine from the protein and NHS on the beads. Primary amine exists at the N terminus of the protein as well as the side chain of lysine residues. There are multiple lysine residues in the Avi tag and SUMO^EU1^ tag at the N terminus of the nanobody, which provide conjugation sites, and the conjugation to these sites possibly ensures the good flexibility and accessibility of the GFP nanobody for autophagosomes binding. We tried to simplify the GFP nanobody by removing the Avi tag and SUMO^EU1^ tag which are not in use and substituting with ten lysine residues separated by linker sequence (GS or GGS) to provide the conjugation sites, however, the autophagosome isolation efficiency was much lower using NHS beads conjugated with this modified nanobody, possibly because that the flexibility of nanobody on the beads was too low to effectively bind to bulky autophagosomes.

### Preparation of SEC (Size Exclusion Chromatography) column

To pack SEC columns, 15 ml (dry matrix volume) of Sepharose 2B-CL matrix (Sigma, CL2B300) was added into an empty gravity column (Bio-Rad, 7321010) with caution to avoid air bubbles trapped in the matrix. A porous 30 μm polyethylene upper bed support was added on top of the matrix to retain the matrix particles. After the liquid flew though, 15 ml (1 CV) 20% ethanol was added to the column, and then another 1 CV 20% ethanol was added before the column was closed for long-term storage at 4°C. Before the usage for autophagosome isolation, the column was washed three times with 1 CV KPBS each time. For each wash, the liquid was allowed to pass through the column completely before new KPBS was added for the next wash. After the three washes, the column is fully equilibrated with KPBS and ready to be used in the isolation experiments. If the column is not used immediately after wash, 1 to 2 ml of KPBS was added to the column after the third wash to prevent the column from drying following which the column was closed and stored at 4°C for short period of time (one or two days). Right before loading the sample (tissue or cell lysate; see the section “autophagosome isolation from tissues and cells”) onto the column, let the KPBS solution completely drain through. After usage, the column should be washed three times (1 CV/wash) with Milli-Q H_2_O to flush off everything from the sample, and then one time with 1 CV 0.5 M NaOH and another three times with Milli-Q H_2_O. At last, the column was washed one time with 1 CV 20% ethanol, and then another 1 CV 20% ethanol was added before the column was closed for long-term storage at 4°C. Notably, the column can be re-used multiple times if properly cleaned after each usage.

### Autophagosome isolation from tissues

For autophagosome isolation from GFP-LC3 mouse skeletal muscles, both quadricep muscles from one mouse were removed and briefly washed in 5 ml cold KPBS. The muscles were then padded dry on tissue paper, and each muscle was put into 900 μl cold isolation buffer (KPBS + proteinase inhibitors+ phosphatase inhibitors) in a 2 ml centrifuge tube. The muscle was diced into small pieces by scissor cutting (70 to 100 times). The buffer containing tissue pieces of both muscles were poured together into a prechilled glass douncer tube (3 ml capacity; to be used with motor-driven pestle later; DWK Life Sciences, 886000-0020). Another 700 µl of cold isolation buffer was used to wash the two centrifuge tubes sequentially to get the remaining tissue pieces, and this washing buffer was also added into the douncer tube (total ∼ 2,500 µl isolation buffer in the tube). The pestle was attached to the motor-drive homogenizer (Glas-Col, 099CK54), and pushed into the douncer tube containing the tissue pieces. With the motor turned on(turning clockwise at a speed ∼2000 rpm), the pestle was slowly pushed to the bottom of the douncer tube before being pulled up till reaching the surface of the liquid (considered as stroke 1). Ten strokes were performed at room temperature during which time the disperse of the tissue can be monitored clearly, then additional 15 strokes were performed on ice. The douncer tube was then put on ice to cool down, meanwhile, any big tissue pieces stuck on the pestle were taken and put back into the douncer tube. Tough tendons tend to be stuck at the bottom of the douncer tube and should be taken out for better homogenization of the tissues. Another 25 strokes were performed on ice and then the lysate was poured evenly into two prechilled 1.5 ml centrifuge tubes. Please note that the homogenization procedure described here is optimized for skeletal muscle, which is hard to break; for other softer tissues, it may not be necessary to perform as many strokes as for skeletal muscle to generate the tissue lysate. After homogenization, the crude tissue lysates in the two centrifuge tubes were centrifuged three times at 700 *g*, 4°C for 5 min each time to get rid of any tissue debris. After the last centrifugation, supernatant from both tubes were poured together to get the final tissue lysate (∼1.5 to 1.7 ml). The tissue lysate was loaded onto the SEC column pre-washed with KPBS as described in the previous section and 4.5 ml void volume was collected. During this time, when the lysate completely passed through, about 7 to 8 ml of cold KPBS were added onto the column. After collecting the void volume, 3 ml elute that contains the majority of mature autophagosomes was collected. Please note that the void volume and elution volume were pre-determined for specifically using 15 ml column matrix and ∼1.5 ml lysate. During the pre-determination process, 0.5 ml fractions were collected sequentially, and the presence of autophagosome or phagophore marker proteins were determined in each fraction by western blots. If the column volume or/and lysate volume changes, the optimal void volume and elution volume will change and need to be determined again. Once the 3 ml elute that contains autophagosomes was obtained, BSA (final concentration 5 μg/μl), proteinase inhibitors and phosphatase inhibitors were added to the elute before 150 ul GFP-nanobody conjugated NHS-activated magnetic beads (the preparation was described in the related section above) were added. The mixture was incubated for 30 min on a rotator at 4°C before the beads were washed five times with cold KPBS. After the wash, beads can be resuspended in the buffer of choice for downstream analysis. For the label-free quantitative Mass Spectrometry in this study, 50 μl of 2×Laemmili buffer was used to resuspend the beads. The beads were then incubated for 15 min at 37°C with occasional mixing before the supernatant was taken for further analysis. In subsequent experiments, we found that incubating the beads with 0.2% Triton X-100 (v/v) in PBS for 10 minutes at 4°C effectively elutes autophagosome contents, while proteins bound directly to GFP-LC3 remain on the beads. This elution method minimizes contamination from non-specific binding proteins when assessing the autophagy cargos but may not recover the autophagy receptors. Therefore, the choice of elution strategy should be determined by the end user based on their specific experimental objectives. Notably, the addition of BSA during isolation step can reduce the non-specific binding of proteins to the beads. For autophagosome isolation during starvation and exercise, quadricep muscles (both) from one GFP-LC3 mouse were sufficient to obtain enough autophagosomes for downstream analysis. For autophagosome isolation during basal condition, quadricep muscles (both) from two GFP-LC3 mice were used, in which case the SEC elute from the two mice were combined together (total 6 ml) before 150 ul GFP-nanobody conjugated beads were added to carry out the later steps as described above. Notably, when performing autophagosome isolation from GFP-LC3 mice, muscles from WT mice subjected to the same treatment are processed at the same time in the same way to be used as non-specific binding controls.

### Trypsin digestion assay

The isolated autophagosomes on the beads from the skeletal muscle of a starved mouse were divided into three portions (50 μl/portion) with one portion left untreated, one portion treated with 2 ug trypsin, and the other portion treated with 2 ug trypsin (Sigma, T4799) with the presence of 0.2% (v/v) Triton X-100. The treatments were carried out for 10 min on ice with occasional mixing before 50 ug trypsin inhibitor (Invitrogen, 17075029) was added into each condition. Laemmili buffer (2×) was added to the samples last, and the samples were boiled immediately at 95 for 10 min before SDS-PAGE and western blots.

### Label-free proteomic and data analysis

Ten percent of the autophagosome preparations (in 2×Laemmili buffer) were run into 4-20% stain-free precast gel (Bio-Rad, 4568096) to estimate the protein concentration in the samples using the stain-free function. Equal amounts of protein from the GFP-LC3 preparations (using the maximum available amount, along with equivalent portions from the corresponding WT controls) or 10 μg total tissue lysates (in 2×Laemmili buffer) were run into 4–20% precast gel (Bio-Rad, 4561094) for around 1 cm distance. The gel was then stained with GelCode™ Blue Stain (Thermo Scientific, 24590) briefly before destained thoroughly with ultrapure water to visualize the areas containing proteins. Those areas were cut-off from the gel and briefly rinsed with 50% acetonitrile in water. The gel slices were subsequently cut into 1×1 mm pieces and subjected to in-gel digestion using trypin as described previously [53].

An aliquot (around a quarter) of the tryptic digest (in 2% acetonitrile/0.1% formic acid in water) was analyzed by LC/MS/MS on an Orbitrap Fusion^TM^ Tribrid^TM^ mass spectrometer (Thermo Scientific) interfaced with a Dionex UltiMate 3000 Binary RSLCnano System. Peptides were separated onto an analytical C18 column (100 μm ID x 25 cm, 5 μm, 18Å Reprosil-Pur C18-AQ beads from Dr Maisch, Ammerbuch-Entringen, Germany) at a flow rate of 350 nl/min. Gradient conditions were: 3-22% B for 90 min; 22-35% B for 10min; 35-90% B for 10 min; 90% B held for 10 min (solvent A, 0.1% formic acid in water; solvent B, 0.1% formic acid in acetonitrile). The peptides were analyzed using data-dependent acquisition method. Orbitrap Fusion was operated with measurement of FTMS1 at resolutions 120,000 FWHM, scan range 350-1500 m/z, AGC target 2E5, and maximum injection time of 50 ms. During a maximum 3 second cycle time, the ITMS2 spectra were collected at rapid scan rate mode, with HCD NCE 34%, 1.6 m/z isolation window, AGC target 1E4, maximum injection time of 35 ms, and dynamic exclusion was employed for 20 seconds.

The raw data files were processed using Thermo Scientific^TM^ Proteome Discoverer^TM^ software version 1.4. Spectra were searched against the Uniprot-mouse database using Sequest. Search results were trimmed to 1% FDR for strict and 5% for relaxed condition using Percolator. For the trypsin, up to two missed cleavages were allowed. MS tolerance was set 10 ppm; MS/MS tolerance 0.6 Da. Carbamidomethylation on cysteine residues was used as fixed modification; oxidation of methione as well as phosphorylation of serine, threonine was set as variable modifications.

Further analysis of the results was done using Microsoft Excel 2019 and GraphPad Prism 10. For each pair of samples (preparations from GFP-LC3 and WT control mice), the proteins whose abundance (the intensity of the corresponding MS spectrum) was at least 1.5 times more in the GFP-LC3 preparation than that in WT control preparation was considered “from autophagosomes”. The common proteins “from autophagosomes” were identified in the four replicates, and only the ones whose abundance are significantly higher in the GFP-LC3 preparations compared to the related WT preparations (*p* ≤ 0.05 determined by paired Student’s t-test) were picked for further analysis. For proteins that were found only in GFP-LC3 samples but not the WT samples, the abundance value was set as 10^4^ (the minimal detection limit) for statistical analysis. The subcellular localization of these proteins was assigned based on the information in Uniprot database. Occasionally, the subcellular localization was further confirmed and modified based on the information obtained via manually searching through literatures. Proteins that are localized to multiple components were counted for each of their respective localizations. To calculate the relative abundance of the proteins in each condition, the original intensity value of each protein was divided by the molecular weight of the protein, and the obtained value was further normalized based on the value of LC3B in the same sample or total protein abundance (refer to the Figure legends). After normalization, the values of all the proteins with the same subcellular localization were added together to reflect the total abundance of proteins from the specific cellular compartment.

### Western blots

All the primary antibodies and secondary antibodies used for western blots were purchased from commercial sources. Anti-GFP antibody (A-6455) was from Invitrogen. Anti-LC3B antibody (NB100-2220) was from Novus Biologicals. Anti-COXIV (11242-1-AP), anti-GAPDH (60004-1-Ig), anti-GM130 (11308-1-AP), anti-Cathepsin D (21327-1-AP), anti-ATP1A2 (16836-1-AP), anti-mouse LAMP1 (65050-1-Ig), anti-FAM134B (21537-1-AP) and anti-TEX264 (25858-1-AP) antibodies were from Proteintech. Anti-p62/SQSTM1 (P0067) and anti-Calnexin (C-terminus, C4731) antibodies were from Sigma. Anti-ATG5 (12994), anti-SEC61A1 (14868), anti-RPS6 (2217), anti-Rab5A (46449) and anti-Rab7 (95746) antibodies were from Cell Signaling Technology. Anti-phospho-ATG16L1 Ser278 (ab195242) was from Abcam. Secondary antibodies used in the study include HRP conjugated anti-rabbit IgG antibody (SA00001-2; for regular western blots), HRP conjugated anti-rabbit IgG (light chain specific) antibody (SA00001-7L; for immunoprecipitation samples) and HRP conjugated anti-mouse IgG antibody (SA00001-1) were from Proteintech. MINI-PROTEAN TGX stain-free precast gels (4-20%; Bio-Rad, 4568096) were used for all the experiments.

### Assessment of mitophagy using Mito-QC mice

Mito-QC mice were subjected to basal, starvation and exercise conditions as described above. After the treatment, mice were euthanized by isoflurane inhalation and perfused with fixation buffer containing 4% paraformaldehyde (PFA) in 100 mM Hepes pH 7.0 (35 ml per mouse at a rate of 4.5 ml/min). After perfusion, quadricep muscles were further fixed in fixation buffer for O/N at 4 before being put sequentially into 15% (w/v) sucrose in 100 mM Hepes pH 7.0 and 30% (w/v) sucrose in 100 mM Hepes pH 7.0 for dehydration. Tissues were then put into Optimal Cutting Temperature compound (OCT) for frozen sectioning. Slides were air-dried and mounted with Prolong Diamond Antifade Mountant with DAPI (Invitrogen). Slides were examined and pictures were taken using an ECHO Revolve microscope. For each mouse, 8 to 10 images were taken randomly. The number of red puncta and the corresponding tissue area in each image were quantified using a Python-based image analysis script. Specifically, we calculated the percentage of the total area (∼160,000 μm²) occupied by the tissue. The analysis script is available for download at https://github.com/slyu13/RFP_Dots_Counting_and_Gap_Area_Detection.git.

### Immunofluorescence

Wild-type mice were subjected to basal, starvation and exercise conditions and frozen sections of the quadricep muscles were prepared as described above. The sections were air-dried and treated with cold acetone (-20) for 20 min in the freezer before being washed three times with PBS (15 min each time). The tissues were then treated with Trueblack Plus Autofluorescence Quencher reagent (Biotium, 23014) according to the user’s manual. The slides were then washed three times with PBG buffer (containing 0.5% [w/v] BSA and 0.2% [w/v] cold water fish gelatin in PBS) and blocked with PBG buffer containing 5% normal donkey serum for 1 hour. Primary antibodies including anti-RPS6 (Cell Signaling Technology, 2217) and anti-mouse LAMP1 (Proteintech, 65050-1-Ig) were diluted in PBG buffer and applied to the tissue sections O/N at 4. Slides were then washed three times with PBG before incubation with secondary antibodies containing donkey anti-rabbit Alexa 647 (Invitrogen, A-31573) and donkey anti-rat Alexa 568 (Invitrogen, A78946) diluted in PBG buffer for 1 hour at room temperature (protected from the light). After washing three times with PBG buffer, the slides were mounted with Prolong Diamond Antifade Mountant with DAPI (Invitrogen). Images were taken using a Nikon AX confocal microscope with tissue area randomly selected. For the tissue from each mouse, 5 to 6 images were taken. The numbers of colocalization dots between RPS6 and LAMP1 were manually counted in 40 to 60 muscle fibers per mouse.

### Co-immunoprecipitation

To determine the interaction between FAM134B or TEX264 with GFP-LC3 II during starvation and exercise, the SEC elute (3 ml) was obtained after the tissue lysate preparation and SEC elution as described above in the section of “Autophagosome isolation from tissues and cells”. BSA (2 μg/μl), proteinase inhibitors and phosphatase inhibitors were added into the elute before 1% Triton X-100 (v/v) was added to release proteins into the solution. For GFP-LC3 II immunoprecipitation, 75 μl single nanobody conjugated magnetic beads were added and the mixture was incubated for 30 min at 4°C before the beads were washed five times with KPBS+1% Triton X-100. For TEX264 immunoprecipitation, 3 μg of anti-TEX264 antibody (Proteintech, 25858-1-AP) and 30 μl pre-washed Protein G Dynabeads were added to the elute and the mixture was incubated for 2 hours at 4°C before the beads were washed five times with KPBS+1% Triton X-100. After the final wash, 50 μl of 2×Laemmili buffer was used to resuspend the beads. The beads were then incubated for 15 min at 55°C with occasional mixing before the supernatant was taken for further analysis.

### Cyro-EM

To visualize the purified autophagosomes using cryo-electron microscopy (cryo-EM), after performing the autophagosome isolation steps from the skeletal muscles of four starved GFP-LC3 mice as described above, the beads containing autophagosomes were combined and eluted twice with 100 μl 0.1 M triethylamine (pH 11.8) with 10 min incubation time (on ice) each time. After each elution, 10 μl 1 M MES (pH 3.0) was added to neutralize the buffer. The eluates were combined and concentrated using Amicon Ultra centrifugal filter unit (3,000 mw cut-off) with the buffer exchanged twice to KPBS. The final eluate (around 50 μl) was used for cryo-EM grid preparation. The sample was vitrified by plunge-freezing in liquid ethane using a FEI Vitrobot Mark IV (ThermoFisher Scientific). Three microliters of the sample were applied onto a glow-discharged Quantifoil copper R 2/1 grid (Quantifoil Micro Tools, GmbH) containing a thin continuous carbon film, and blotted for 2 s at 100% humidity at room temperature. The grids were examined in liquid nitrogen using a Glacios Microscope (ThermoFisher Scientific) operated at 200 Kv. The images were acquired using Falcon 4i Camera with 10 EV energy filter (Selectris X) at a magnification of ×130,000 with a calibrated physical pixel size of 0.92 Å.

### Data analysis

Graphs were made using GraphPad Prism 10. Statistical analysis to compare the differences between conditions were performed using two-sides, unpaired Student’s t-test either in Microsoft Excel or GraphPad Prism 10.

### Data Availability

The mass spectrometry data from this study have been deposited to the Open Science Framework database [https://osf.io/gf5wd/?view_only=d10bfe8438984bafa4e3c7909b2ecc30]. All the data are available on reasonable request to the corresponding author. There is a preprint version of this manuscript on bioRxiv [54].

### Usage of Generative Artificial Intelligence (AI)

We used OpenAI’s ChatGPT (GPT-4, accessed via ChatGPT Plus) to assist with improving the clarity and flow of the manuscript during the writing process. All scientific content, data interpretation, and final revisions were performed and approved by the authors.

## Authors contributions

MF-investigation, data curation; SL-investigation, methodology, data curation; TPN-investigation, data curation; DZ-investigation, data curation; HL-investigation, data curation; YZ-methodology, data curation; YAA-conceptualization, methodology; HY-conceptualization, methodology; GD-conceptualization, methodology; YL-conceptualization, methodology, writing, funding acquisition.

## Acknowledgements

This work is supported in part by the Clinical and Translational Proteomics Service Center and the Cryo-EM core facility at the University of Texas Health Science Center at Houston. We also thank Proteomic Core at the University of Texas Southwestern Medical Center for their technical support.

## Disclosure and competing interests

The authors declare that they have no conflict of interest.

## Funding resources

This work is supported by the funding from the McGovern Medical School, University of Texas Health Science Center at Houston.

## Abbreviation

ER: endoplasmic reticulum
GABARAP: Gamma-aminobutyric acid receptor-associated protein
LC3: Microtubule-associated proteins 1A/1B light chain 3
SEC: size exclusion chromatography
SR: sarcoplasmic reticulum
WT: wild-type.

---

**Figure S1.**
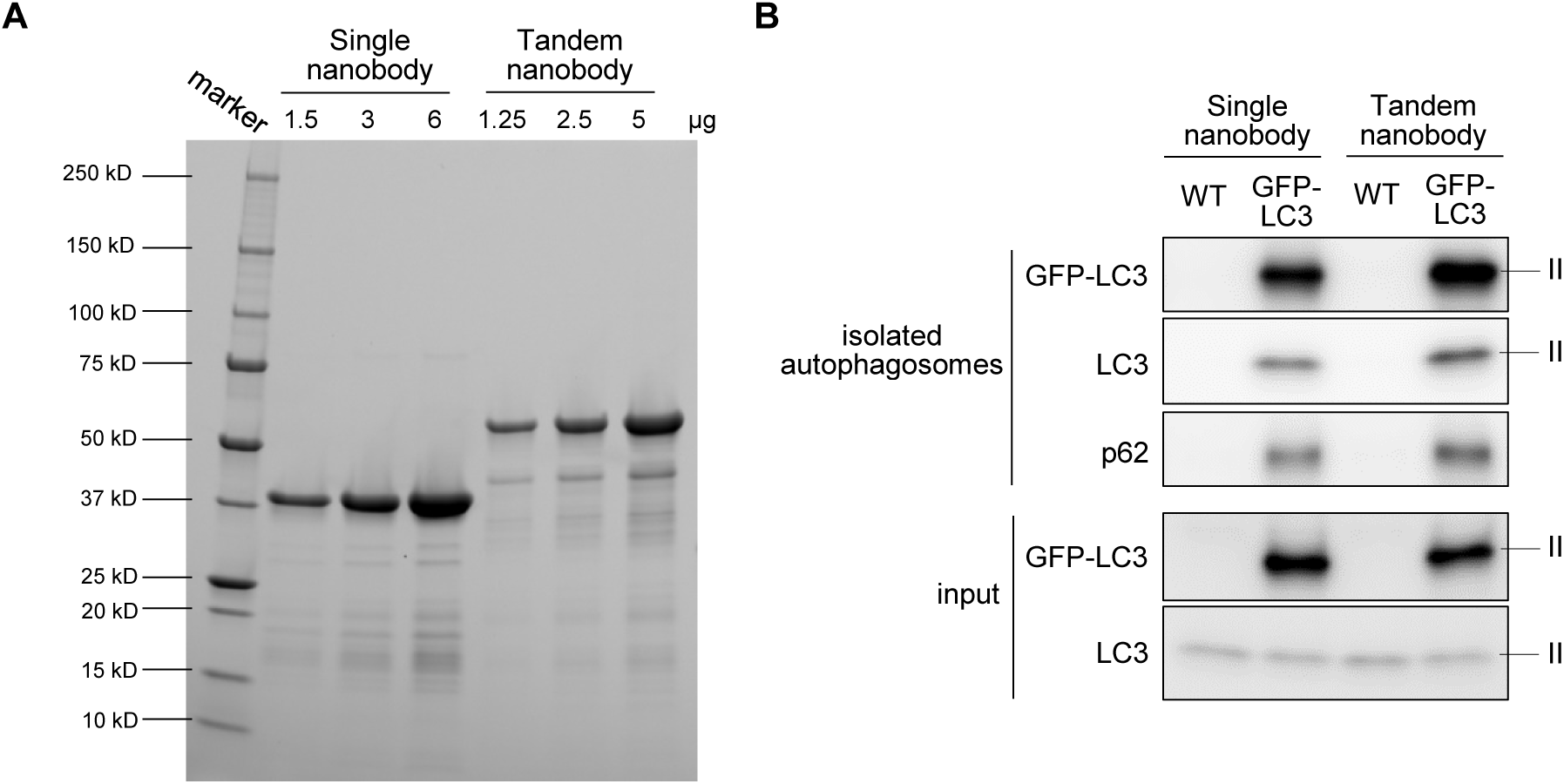
Comparing the autophagosome isolation efficiency using tandem GFP nanobody and single GFP nanobody. (**A**) Purified single GFP nanobody and tandem GFP nanobody detected in the stain-free gel. Different amounts of purified proteins were loaded. (**B**) Western blot analysis shows increased enrichment of GFP-LC3 II, LC3 II, and p62 in autophagosome preparations from skeletal muscles of starved GFP-LC3 mice when using tandem GFP nanobody-conjugated beads compared to single GFP nanobody-conjugated beads during the pull-down step. Similar results were observed in at least three independent experiments.

**Figure S2.**
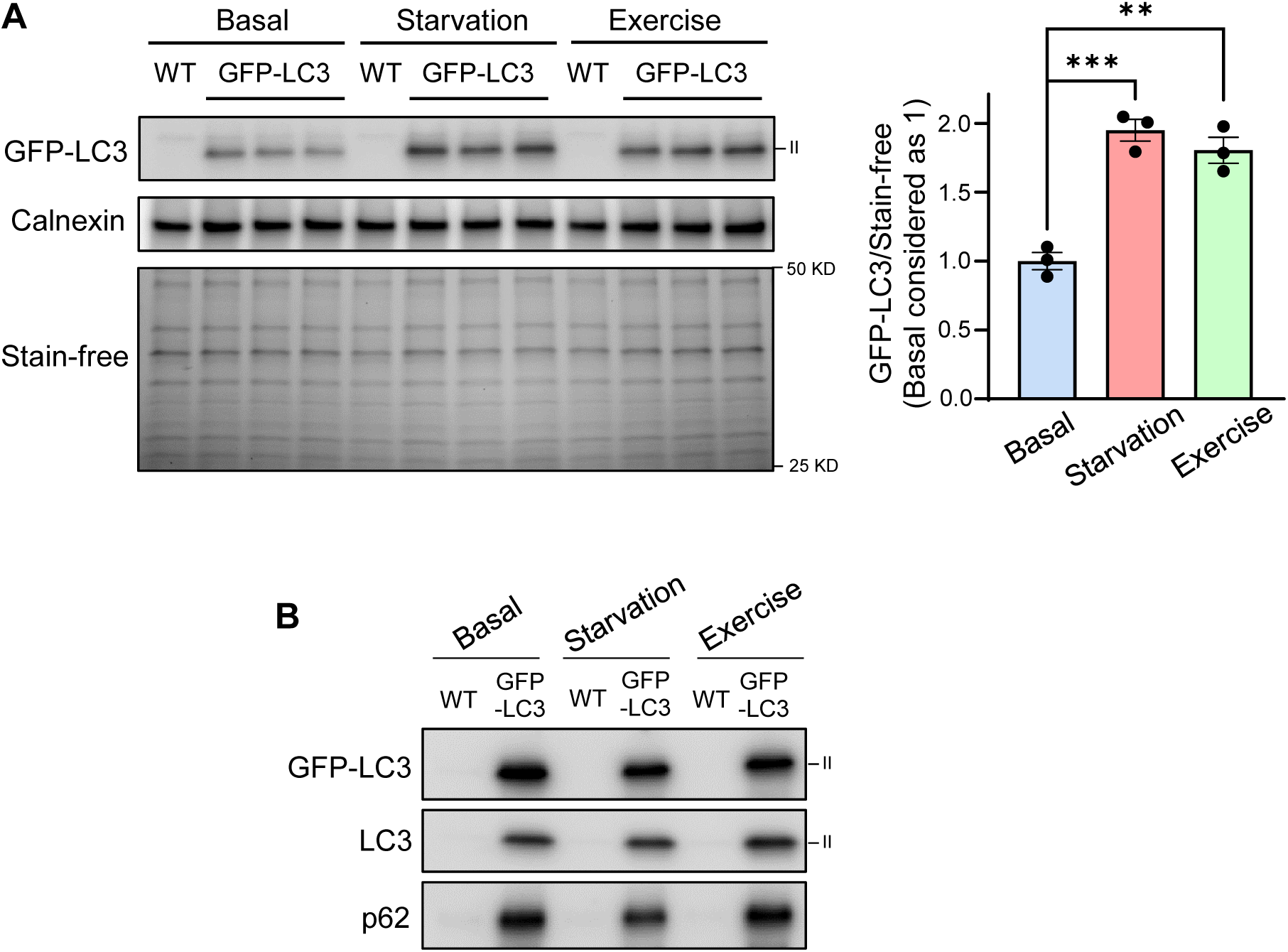
Autophagosome isolation from the skeletal muscles of GFP-LC3 mice under basal, starvation and exercise conditions. (**A**) Left, western blot analysis shows the levels of lipidated GFP-LC3 II in skeletal muscles lysates from WT and GFP-LC3 mice (post SEC to eliminate GFP-LC3 I) under basal, starvation and exercise conditions. Calnexin and total protein levels determined by the staining signals from the stain-free gel were used as the loading controls. Right, quantification of GFP-LC3 II levels normalized to the stain-free signals as shown on the left (n=3). (**B**) Western blot analysis shows the enrichment of GFP-LC3 II, LC3 II, and p62 in the autophagosome preparations from the skeletal muscles of GFP-LC3 mice compared to those from WT mice under basal, starvation and exercise conditions. ** *p* <0.01, *** *p* < 0.001 when compared to basal condition. Two sides, unpaired Student’s t-test was used for statistical analysis. Data are mean ± s.e.m.

**Figure S3.**
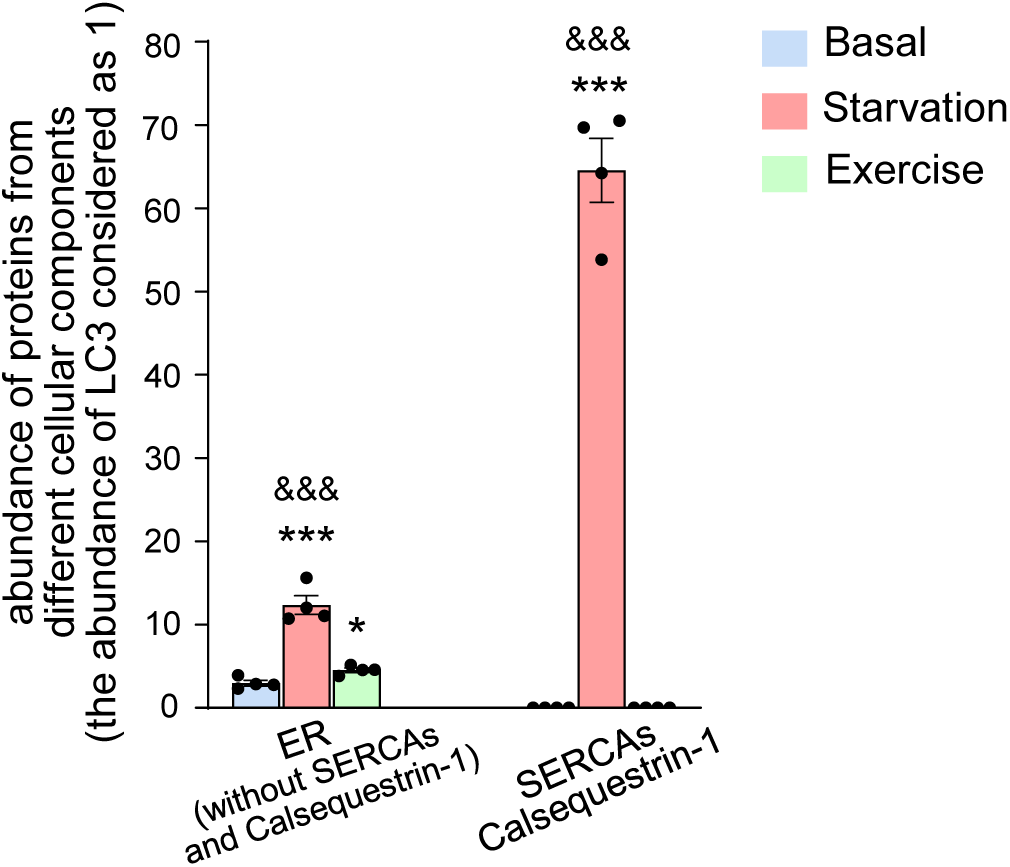
Autophagosomes from the skeletal muscle under starvation condition contain a higher abundance of ER proteins compared to those obtained under basal or exercise condition. On the right side of the graph, the abundance of SERCAs and Calsequestrin-1 identified in the isolated autophagosomes under basal, starvation and exercise conditions are shown. Notably, these proteins were not identified under basal nor exercise condition, thus the abundance values are zero. On the left side of the graph, the abundance of all ER proteins, excluding SERCAs and Calsequestrin-1, identified in the isolated autophagosomes under basal, starvation and exercise conditions is shown. The abundance of proteins is normalized to the abundance of LC3B protein in each replicate. * *p* < 0.05, *** *p* < 0.001 when compared to basal condition. $$$ *p* < 0.001 when compared to exercise condition. Two sides, unpaired Student’s t-test was used for statistical analysis. Data are mean ± s.e.m.

**Figure S4.**
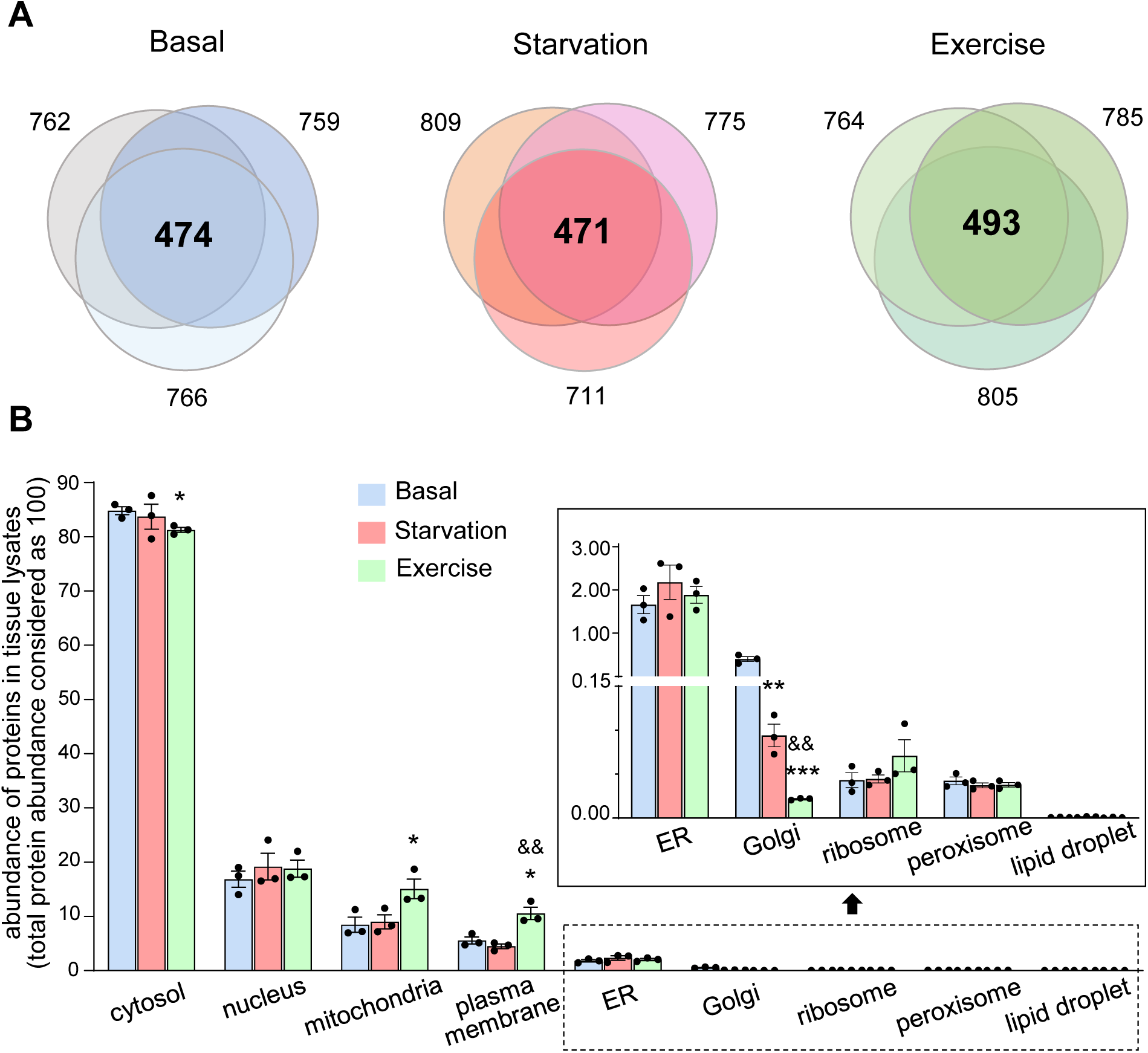
The abundance of proteins from various cellular components in the tissue lysates of skeletal muscle under basal, starvation and exercise conditions. (**A**) The identification of proteins in the tissue lysates of skeletal muscle under basal, starvation and exercise conditions. Each circle stands for one replicate, and the number near it indicates how many proteins were identified in this replicate. The number in the overlapping region is the number of proteins identified in all three replicates. These proteins were subjected to further analysis in (B). (**B**) The total abundance of proteins identified in (A) from various cellular components under basal, starvation and exercise conditions. The total protein abundance in each replicate was considered as 100. Proteins that are localized to multiple components were counted for each of their respective localizations. * *p* < 0.05, ** *p* <0.01, *** *p* < 0.001 when compared to basal condition. $$ *p* < 0.01 when compared to starvation condition. Two sides, unpaired Student’s t-test was used for statistical analysis. Data are mean ± s.e.m.

**Table S1.** Peptides of RTN3 (Uniprot ID: Q9ES97) identified with Mass spectrometry.

| Peptide sequence identified | Corresponding position in RTN3 |
| --- | --- |
| TQIDHYVGIAR | amino acids 931-941 |
| SEEGHPFK | amino acids 838-845 |
| SVIQAVQK | amino acids 830-837 |

## Notes

https://osf.io/gf5wd/?view_only=d10bfe8438984bafa4e3c7909b2ecc30

### Competing Interest Statement

The authors have declared no competing interest.

### Summary of Updates

main text of the manuscript updated; Figures revised; Authorship changed

